## Supplemental information for "Instantly adhesive and ultra-elastic patches for dynamic organ and wound repair"

**This PDF file includes:**

Figs. S1 and S2

Tables S1 to S3

**Other Supplementary Materials for this manuscript include the following:**

Movies S1 to S9

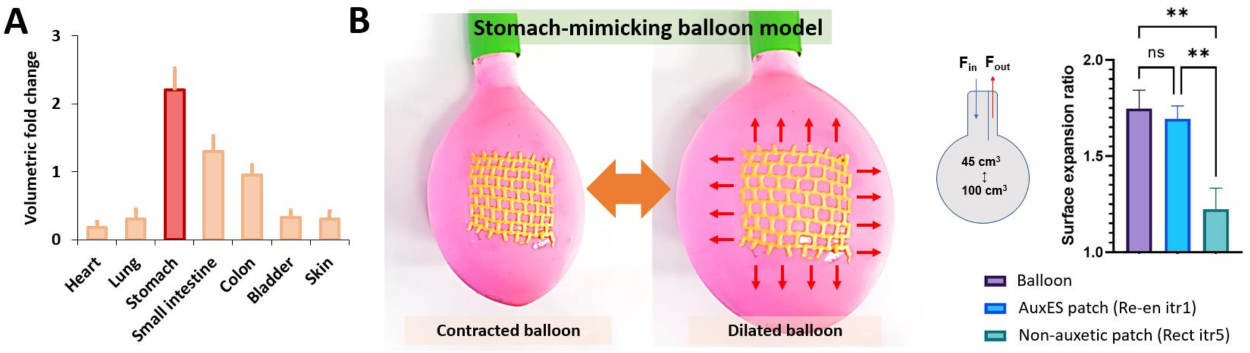


**Figure S1. A.** Volumetric fold change of different dynamic organs. The stomach has the highest volumetric fold change compared with other dynamic organs*(1)*. **B.** The AuxES patch (Re-en itr1 architecture) tested on a dilating balloon model demonstrated adherence to the balloon mechanics and a similar surface expansion to that of the balloon (see **Supplementary Video S4**), while the non-auxetic patch (Rect itr5) did not conform to the deformation of the balloon. **represents $p < 0.01$ (group-wise comparisons carried by Tukey's HSD post-hoc tests).

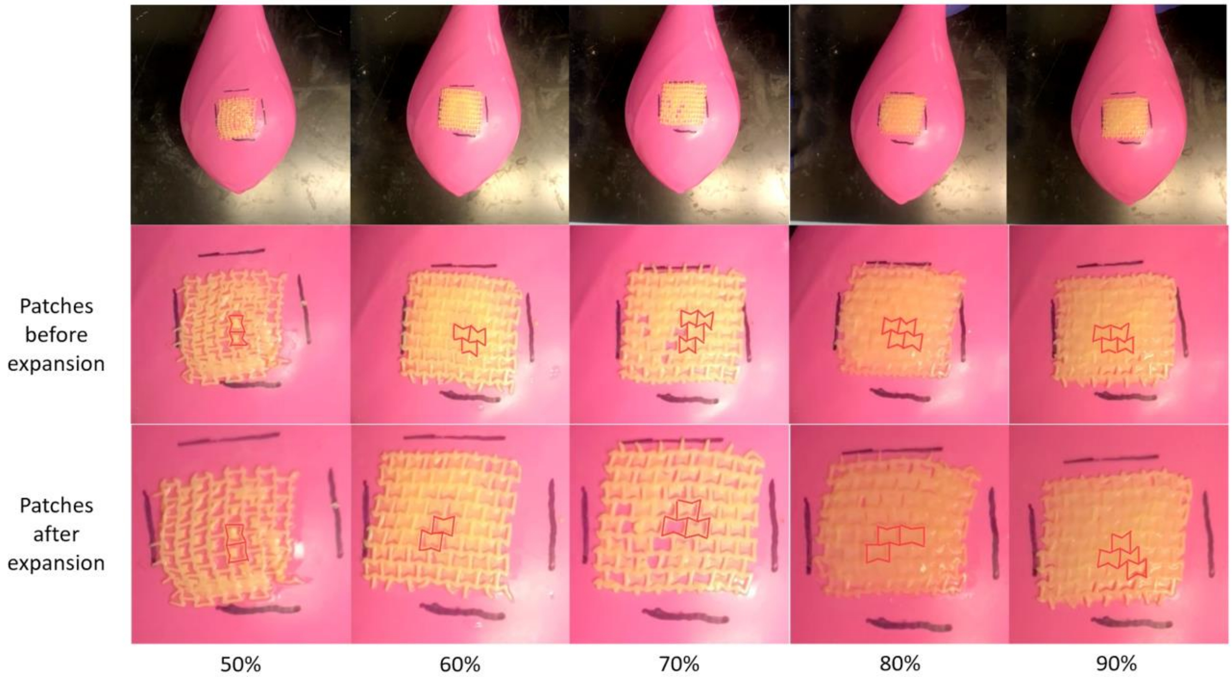


**Figure S2.** Different compositions of the hole-filling material were fabricated under different UV exposure durations and tested for their compliance to the balloon model. The numbers 50-90% at the bottom denote the ink composition as per Table S2. With 15 s crosslinking time, 50% and 60% ink formulation (as per Table S2) cannot crosslink with the original matrix, i.e., the holes cannot be sealed. 70% successfully crosslinked and maintained the auxetic properties, but the hole-filling material was prone to detachment. The ink composition with concentration of the hole filling material was 80% v/v as that of the lattice material demonstrated void-filling capability while also exhibiting auxetic characteristics. 90% ink started to cause the loss of auxetic property and 100% ink resulted in complete loss of the auxetic property.

**Table S1.** Dimensions of the design elements of the different patch architectures.

| Iteration | Geometry | H (mm) | w (mm) | t (mm) | ia ($^0$) | r (mm) | te (mm) |
|---|---|---|---|---|---|---|---|
| 1 | Re-entrant honeycomb | 3 | 3 | 0.75 | 90 | N/A | N/A |
| 2 | Re-entrant honeycomb | 2.25 | 2.25 | 0.75 | 90 | N/A | N/A |
| 3 | Re-entrant honeycomb | 1.5 | 1.5 | 0.75 | 90 | N/A | N/A |
| 4 | Re-entrant honeycomb | 3 | 2 | 0.75 | 90 | N/A | N/A |
| 5 | Re-entrant honeycomb | 2.25 | 1.5 | 0.75 | 90 | N/A | N/A |
| 6 | Re-entrant honeycomb | 1.5 | 1 | 0.75 | 90 | N/A | N/A |
| 7 | Re-entrant honeycomb | 3 | 3 | 1.125 | 90 | N/A | N/A |
| 8 | Re-entrant honeycomb | 2.25 | 2.25 | 1.125 | 90 | N/A | N/A |
| 9 | Re-entrant honeycomb | 1.5 | 1.5 | 1.125 | 90 | N/A | N/A |
| 10 | Re-entrant honeycomb | 3 | 2 | 1.125 | 90 | N/A | N/A |
| 11 | Re-entrant honeycomb | 2.25 | 1.5 | 1.125 | 90 | N/A | N/A |
| 12 | Re-entrant honeycomb | 1.5 | 1 | 1.125 | 90 | N/A | N/A |
| 13 | Re-entrant honeycomb | 3 | 3 | 0.75 | 112.5 | N/A | N/A |
| 14 | Re-entrant honeycomb | 2.25 | 2.25 | 0.75 | 112.5 | N/A | N/A |
| 15 | Re-entrant honeycomb | 1.5 | 1.5 | 0.75 | 112.5 | N/A | N/A |
| 16 | Re-entrant honeycomb | 3 | 2 | 0.75 | 112.5 | N/A | N/A |
| 17 | Re-entrant honeycomb | 2.25 | 1.5 | 0.75 | 112.5 | N/A | N/A |
| 18 | Re-entrant honeycomb | 1.5 | 1 | 0.75 | 112.5 | N/A | N/A |
| 19 | Re-entrant honeycomb | 3 | 3 | 1.125 | 112.5 | N/A | N/A |
| 20 | Re-entrant honeycomb | 2.25 | 2.25 | 1.125 | 112.5 | N/A | N/A |
| 21 | Re-entrant honeycomb | 1.5 | 1.5 | 1.125 | 112.5 | N/A | N/A |
| 22 | Re-entrant honeycomb | 3 | 2 | 1.125 | 112.5 | N/A | N/A |
| 23 | Re-entrant honeycomb | 2.25 | 1.5 | 1.125 | 112.5 | N/A | N/A |
| 24 | Re-entrant honeycomb | 1.5 | 1 | 1.125 | 112.5 | N/A | N/A |
| 1 | Lozenge truss | 2.25 | N/A | 0.375 | N/A | N/A | 0.75 |
| 2 | Lozenge truss | 1.5 | N/A | 0.375 | N/A | N/A | 0.75 |
| 3 | Lozenge truss | 3 | N/A | 0.375 | N/A | N/A | 0.75 |
| 4 | Lozenge truss | 3 | N/A | 0.75 | N/A | N/A | 0.75 |
| 5 | Lozenge truss | 1.5 | N/A | 0.75 | N/A | N/A | 0.75 |
| 6 | Lozenge truss | 2.25 | N/A | 0.75 | N/A | N/A | 0.75 |
| 7 | Lozenge truss | 3 | N/A | 0.375 | N/A | N/A | 1.125 |
| 8 | Lozenge truss | 2.25 | N/A | 0.75 | N/A | N/A | 1.125 |
| 1 | Sinusoidal ligaments | 3 | 3 | 0.75 | N/A | 1.875 | N/A |
| 2 | Sinusoidal ligaments | 2.25 | 2.25 | 0.75 | N/A | 1.875 | N/A |
| 3 | Sinusoidal ligaments | 1.5 | 1.5 | 0.75 | N/A | 1.875 | N/A |
| 4 | Sinusoidal ligaments | 3 | 2 | 0.75 | N/A | 1.875 | N/A |
| 5 | Sinusoidal ligaments | 2.25 | 1.5 | 0.75 | N/A | 1.875 | N/A |
| 6 | Sinusoidal ligaments | 1.5 | 1 | 0.75 | N/A | 1.875 | N/A |
| 7 | Sinusoidal ligaments | 3 | 3 | 1.125 | N/A | 1.875 | N/A |

| | | | | | | | |
|---|---|---|---|---|---|---|---|
| 8 | Sinusoidal ligaments | 2.25 | 2.25 | 1.125 | N/A | 1.875 | N/A |
| 9 | Sinusoidal ligaments | 3 | 2 | 1.125 | N/A | 1.875 | N/A |
| 10 | Sinusoidal ligaments | 2.25 | 1.5 | 1.125 | N/A | 1.875 | N/A |
| 11 | Sinusoidal ligaments | 1.5 | 1 | 1.125 | N/A | 1.875 | N/A |
| 12 | Sinusoidal ligaments | 3 | 3 | 0.75 | N/A | 2.25 | N/A |
| 13 | Sinusoidal ligaments | 2.25 | 2.25 | 0.75 | N/A | 2.25 | N/A |
| 14 | Sinusoidal ligaments | 1.5 | 1.5 | 0.75 | N/A | 2.25 | N/A |
| 15 | Sinusoidal ligaments | 3 | 2 | 0.75 | N/A | 2.25 | N/A |
| 16 | Sinusoidal ligaments | 2.25 | 1.5 | 0.75 | N/A | 2.25 | N/A |
| 17 | Sinusoidal ligaments | 3 | 3 | 1.125 | N/A | 2.25 | N/A |
| 18 | Sinusoidal ligaments | 2.25 | 2.25 | 1.125 | N/A | 2.25 | N/A |
| 19 | Sinusoidal ligaments | 3 | 2 | 1.125 | N/A | 2.25 | N/A |
| 20 | Sinusoidal ligaments | 2.25 | 1.5 | 1.125 | N/A | 2.25 | N/A |
| 21 | Sinusoidal ligaments | 1.5 | 1 | 1.125 | N/A | 2.25 | N/A |
| 1 | Honeycomb | 3 | N/A | 0.75 | N/A | N/A | N/A |
| 2 | Honeycomb | 2.25 | N/A | 0.75 | N/A | N/A | N/A |
| 3 | Honeycomb | 3 | N/A | 1.125 | N/A | N/A | N/A |
| 4 | Honeycomb | 2.25 | N/A | 1.125 | N/A | N/A | N/A |
| 1 | Inclined Truss | 3 | 3 | 0.75 | N/A | N/A | N/A |
| 2 | Inclined Truss | 2.25 | 2.25 | 0.75 | N/A | N/A | N/A |
| 3 | Inclined Truss | 1.5 | 1.5 | 0.75 | N/A | N/A | N/A |
| 4 | Inclined Truss | 3 | 3 | 1.125 | N/A | N/A | N/A |
| 5 | Inclined Truss | 2.25 | 2.25 | 1.125 | N/A | N/A | N/A |
| 6 | Inclined Truss | 1.5 | 1.5 | 1.125 | N/A | N/A | N/A |

**Table S2.** Different compositions of the void-filling material (LAP concentration was the same at 0.03% w/v). In this formulation, the 80% v/v GelMA – 20% v/v ACA mixture (termed as the original ink composition) used for the patch lattices was further diluted to 50, 60, 70, 80, and 90% of its original concentration by mixing with PBS and application to the patch voids followed by UV exposure.

| **Composition** | **Amount of Original ink formulation (% v/v) in 1x PBS** | **GelMA (μL)** | **ACA (μL)** | **1x PBS (μL)** |
|---|---|---|---|---|
| 1 | 100% | 500 | 100 | 0 |
| 2 | 90% | 450 | 90 | 60 |
| 3 | 80% | 400 | 80 | 120 |
| 4 | 70% | 350 | 70 | 180 |
| 5 | 60% | 300 | 60 | 240 |
| 6 | 50% | 250 | 50 | 300 |

**Table S3.** Statistical data of Nanostring analysis.

| Gene | Discovery | P value | Mean of No Patch | Mean of EXOs + Curcumin | Difference | SE of difference | t ratio | df | q value |
|---|---|---|---|---|---|---|---|---|---|
| C2 | No | 0.310737 | 1.000 | 2.129 | -1.129 | 0.9741 | 1.160 | 4.000 | 0.459692 |
| C8G | No | 0.888779 | 1.000 | 1.079 | -0.07923 | 0.5318 | 0.1490 | 4.000 | 0.897667 |
| C9 | No | 0.474118 | 1.000 | 2.043 | -1.043 | 1.321 | 0.7892 | 4.000 | 0.498207 |
| CAMP | No | 0.054254 | 1.000 | 0.5100 | 0.4900 | 0.1817 | 2.697 | 4.000 | 0.248196 |
| CCR2 | No | 0.043413 | 1.000 | 2.518 | -1.518 | 0.5206 | 2.916 | 4.000 | 0.225810 |
| CCR3 | No | 0.322673 | 1.000 | 2.999 | -1.999 | 1.774 | 1.127 | 4.000 | 0.459692 |
| CCR4 | No | 0.300292 | 1.000 | 2.283 | -1.283 | 1.079 | 1.189 | 4.000 | 0.459692 |
| GPR29 | No | 0.084662 | 1.000 | 2.814 | -1.814 | 0.7952 | 2.281 | 4.000 | 0.307221 |
| ACKR4 | No | 0.246783 | 1.000 | 3.111 | -2.111 | 1.557 | 1.355 | 4.000 | 0.459692 |
| CCRL2 | No | 0.667334 | 1.000 | 1.365 | -0.3647 | 0.7873 | 0.4632 | 4.000 | 0.680615 |
| Cd1d1 | No | 0.059582 | 1.000 | 2.878 | -1.878 | 0.7202 | 2.607 | 4.000 | 0.258261 |
| CD2 | No | 0.079901 | 1.000 | 2.110 | -1.110 | 0.4757 | 2.334 | 4.000 | 0.307221 |
| CD3E | No | 0.352579 | 1.000 | 2.584 | -1.584 | 1.507 | 1.051 | 4.000 | 0.459692 |
| CD7 | No | 0.170373 | 1.000 | 3.056 | -2.056 | 1.232 | 1.669 | 4.000 | 0.411740 |
| CFB | No | 0.406533 | 1.000 | 1.688 | -0.6875 | 0.7419 | 0.9267 | 4.000 | 0.459692 |
| CFD | No | 0.436346 | 1.000 | 2.265 | -1.265 | 1.464 | 0.8639 | 4.000 | 0.472844 |
| CFH° | No | 0.027294 | 1.000 | 2.427 | -1.427 | 0.4198 | 3.399 | 4.000 | 0.177465 |
| CFI | No | 0.354039 | 1.000 | 2.775 | -1.775 | 1.695 | 1.047 | 4.000 | 0.459692 |
| CIITA° | No | 0.038305 | 1.000 | 2.220 | -1.220 | 0.4008 | 3.043 | 4.000 | 0.217268 |
| CLU | No | 0.050639 | 1.000 | 2.358 | -1.358 | 0.4913 | 2.764 | 4.000 | 0.248196 |
| CR2 | No | 0.262282 | 1.000 | 2.359 | -1.359 | 1.043 | 1.304 | 4.000 | 0.459692 |
| CSF2 | No | 0.257003 | 1.000 | 2.713 | -1.713 | 1.297 | 1.321 | 4.000 | 0.459692 |

| Gene | Discovery | P value | Mean of No Patch | Mean of EXOs + Curcumin | Difference | SE of difference | t ratio | df | q value |
|---|---|---|---|---|---|---|---|---|---|
| CTLA4 | No | 0.381543 | 1.000 | 2.653 | -1.653 | 1.683 | 0.9824 | 4.000 | 0.459692 |
| Cxcl15 | No | 0.384275 | 1.000 | 2.743 | -1.743 | 1.786 | 0.9761 | 4.000 | 0.459692 |
| CXCR1 | No | 0.355534 | 1.000 | 2.514 | -1.514 | 1.451 | 1.044 | 4.000 | 0.459692 |
| DEFB1 | No | 0.078204 | 1.000 | 2.018 | -1.018 | 0.4325 | 2.354 | 4.000 | 0.307221 |
| GATA3* | No | 0.022575 | 1.000 | 2.449 | -1.449 | 0.4016 | 3.609 | 4.000 | 0.174112 |
| GZMB | No | 0.260967 | 1.000 | 2.520 | -1.520 | 1.162 | 1.308 | 4.000 | 0.459692 |
| ICOS | No | 0.412258 | 1.000 | 2.118 | -1.118 | 1.222 | 0.9144 | 4.000 | 0.461153 |
| IFNB1 | No | 0.284856 | 1.000 | 3.958 | -2.958 | 2.398 | 1.234 | 4.000 | 0.459692 |
| IL17B | No | 0.159041 | 1.000 | 3.300 | -2.300 | 1.331 | 1.728 | 4.000 | 0.393930 |
| IL17F | No | 0.293902 | 1.000 | 2.720 | -1.720 | 1.425 | 1.207 | 4.000 | 0.459692 |
| IL18 | No | 0.139785 | 1.000 | 2.785 | -1.785 | 0.9707 | 1.839 | 4.000 | 0.368184 |
| IL2 | No | 0.210897 | 1.000 | 3.110 | -2.110 | 1.418 | 1.488 | 4.000 | 0.459692 |
| IL21 | No | 0.405717 | 1.000 | 2.496 | -1.496 | 1.611 | 0.9285 | 4.000 | 0.459692 |
| IL23R | No | 0.349195 | 1.000 | 2.003 | -1.003 | 0.9467 | 1.059 | 4.000 | 0.459692 |
| IL2RB° | No | 0.005319 | 1.000 | 2.251 | -1.251 | 0.2274 | 5.503 | 4.000 | 0.129937 |
| IL5 | No | 0.174148 | 1.000 | 2.874 | -1.874 | 1.136 | 1.651 | 4.000 | 0.411740 |
| ILF3 | No | 0.158883 | 1.000 | 2.788 | -1.788 | 1.034 | 1.729 | 4.000 | 0.393930 |
| KIR3DL1 | No | 0.362668 | 1.000 | 0.4170 | 0.5830 | 0.5680 | 1.027 | 4.000 | 0.459692 |
| KIR3DL2 | No | 0.447744 | 1.000 | 2.144 | -1.144 | 1.360 | 0.8409 | 4.000 | 0.475294 |
| KLRD1 | No | 0.097358 | 1.000 | 2.505 | -1.505 | 0.6982 | 2.156 | 4.000 | 0.316505 |
| Klra4 | No | 0.207935 | 1.000 | 3.344 | -2.344 | 1.562 | 1.500 | 4.000 | 0.459692 |
| Klra6 | No | 0.290227 | 1.000 | 3.956 | -2.956 | 2.428 | 1.218 | 4.000 | 0.459692 |

| Gene | Discovery | P value | Mean of No Patch | Mean of EXOs + Curcumin | Difference | SE of difference | t ratio | df | q value |
|---|---|---|---|---|---|---|---|---|---|
| Klra7 | No | 0.340173 | 1.000 | 2.774 | -1.774 | 1.640 | 1.082 | 4.000 | 0.459692 |
| Klra8** | No | 0.000105 | 1.000 | 1.936 | -0.9358 | 0.06102 | 15.34 | 4.000 | 0.010969 |
| Klrd1 | No | 0.132000 | 1.000 | 2.532 | -1.532 | 0.8114 | 1.888 | 4.000 | 0.368184 |
| LAIR1 | No | 0.648755 | 1.000 | 1.299 | -0.2995 | 0.6092 | 0.4916 | 4.000 | 0.668217 |
| LILRA6 | No | 0.243468 | 1.000 | 2.216 | -1.216 | 0.8893 | 1.367 | 4.000 | 0.459692 |
| MASP2 | No | 0.445766 | 1.000 | 2.132 | -1.132 | 1.340 | 0.8449 | 4.000 | 0.475294 |
| MBL2 | No | 0.360758 | 1.000 | 2.500 | -1.500 | 1.454 | 1.031 | 4.000 | 0.459692 |
| MUC1 | No | 0.097352 | 1.000 | 0.4594 | 0.5406 | 0.2508 | 2.156 | 4.000 | 0.316505 |
| MX1 | No | 0.398248 | 1.000 | 3.171 | -2.171 | 2.297 | 0.9448 | 4.000 | 0.459692 |
| NFATC2* | No | 0.012490 | 1.000 | 2.159 | -1.159 | 0.2686 | 4.316 | 4.000 | 0.129937 |
| NFKBIZ | No | 0.138045 | 1.000 | 0.4918 | 0.5082 | 0.2748 | 1.850 | 4.000 | 0.368184 |
| NOTCH1* | No | 0.012177 | 1.000 | 2.097 | -1.097 | 0.2524 | 4.348 | 4.000 | 0.129937 |
| NOX3 | No | 0.431643 | 1.000 | 2.101 | -1.101 | 1.261 | 0.8736 | 4.000 | 0.472672 |
| Pdcd1lg2 | No | 0.405976 | 1.000 | 2.347 | -1.347 | 1.452 | 0.9279 | 4.000 | 0.459692 |
| PDCD1LG2 | No | 0.114492 | 1.000 | 3.148 | -2.148 | 1.067 | 2.012 | 4.000 | 0.350313 |
| SELPLG* | No | 0.010164 | 1.000 | 2.588 | -1.588 | 0.3466 | 4.583 | 4.000 | 0.129937 |
| STAT2 | No | 0.398504 | 1.000 | 1.409 | -0.4086 | 0.4327 | 0.9443 | 4.000 | 0.459692 |
| STAT6* | No | 0.010581 | 1.000 | 2.006 | -1.006 | 0.2220 | 4.529 | 4.000 | 0.129937 |
| TCF7 | No | 0.110819 | 1.000 | 2.717 | -1.717 | 0.8411 | 2.041 | 4.000 | 0.349349 |
| TNFRSF11A | No | 0.335217 | 1.000 | 2.823 | -1.823 | 1.665 | 1.095 | 4.000 | 0.459692 |
| TNFRSF17 | No | 0.395986 | 1.000 | 2.470 | -1.470 | 1.547 | 0.9498 | 4.000 | 0.459692 |
| TNFSF11* | No | 0.039682 | 1.000 | 2.275 | -1.275 | 0.4239 | 3.007 | 4.000 | 0.217268 |

| Gene | Discovery | P value | Mean of No Patch | Mean of EXOs + Curcumin | Difference | SE of difference | t ratio | df | q value |
|---|---|---|---|---|---|---|---|---|---|
| TNFSF15 | No | 0.361226 | 1.000 | 2.343 | -1.343 | 1.304 | 1.030 | 4.000 | 0.459692 |
| TNFSF18 | No | 0.381904 | 1.000 | 2.260 | -1.260 | 1.283 | 0.9815 | 4.000 | 0.459692 |
| XCR1 | No | 0.141568 | 1.000 | 2.260 | -1.260 | 0.6892 | 1.828 | 4.000 | 0.368184 |
| CASP2* | No | 0.024736 | 1.000 | 2.658 | -1.658 | 0.4729 | 3.507 | 4.000 | 0.174112 |
| CCL19 | No | 0.141456 | 1.000 | 2.111 | -1.111 | 0.6073 | 1.829 | 4.000 | 0.368184 |
| CD19 | No | 0.294251 | 1.000 | 3.258 | -2.258 | 1.872 | 1.206 | 4.000 | 0.459692 |
| CD24 | No | 0.003966 | 1.000 | 2.780 | -1.780 | 0.2984 | 5.965 | 4.000 | 0.129937 |
| CD55 | No | 0.085880 | 1.000 | 1.869 | -0.8688 | 0.3830 | 2.268 | 4.000 | 0.307221 |
| CD79A | No | 0.257877 | 1.000 | 2.961 | -1.961 | 1.488 | 1.318 | 4.000 | 0.459692 |
| CX3CL1* | No | 0.024928 | 1.000 | 2.087 | -1.087 | 0.3108 | 3.499 | 4.000 | 0.174112 |
| CXCL13 | No | 0.123192 | 1.000 | 3.396 | -2.396 | 1.230 | 1.948 | 4.000 | 0.366162 |
| CXCR6 | No | 0.238941 | 1.000 | 2.132 | -1.132 | 0.8184 | 1.383 | 4.000 | 0.459692 |
| GFI1 | No | 0.500916 | 1.000 | 1.769 | -0.7690 | 1.041 | 0.7390 | 4.000 | 0.521103 |
| IFNG | No | 0.309422 | 1.000 | 2.088 | -1.088 | 0.9354 | 1.163 | 4.000 | 0.459692 |
| IL10 | No | 0.372960 | 1.000 | 2.195 | -1.195 | 1.193 | 1.002 | 4.000 | 0.459692 |
| IL11 | No | 0.054874 | 1.000 | 2.307 | -1.307 | 0.4866 | 2.686 | 4.000 | 0.248196 |
| MAP4K1 | No | 0.246877 | 1.000 | 2.333 | -1.333 | 0.9834 | 1.355 | 4.000 | 0.459692 |
| MAPK11 | No | 0.270403 | 1.000 | 1.903 | -0.9030 | 0.7066 | 1.278 | 4.000 | 0.459692 |
| MS4A1 | No | 0.431117 | 1.000 | 2.055 | -1.055 | 1.206 | 0.8747 | 4.000 | 0.472672 |
| Plau* | No | 0.025105 | 1.000 | 2.035 | -1.035 | 0.2966 | 3.491 | 4.000 | 0.174112 |
| Prdm1** | No | 0.008173 | 1.000 | 2.798 | -1.798 | 0.3686 | 4.878 | 4.000 | 0.129937 |
| RAE1 | No | 0.088596 | 1.000 | 2.139 | -1.139 | 0.5086 | 2.240 | 4.000 | 0.307221 |

| Gene | Discovery | P value | Mean of No Patch | Mean of EXOs + Curcumin | Difference | SE of difference | t ratio | df | q value |
|---|---|---|---|---|---|---|---|---|---|
| TYK2 | No | 0.258352 | 1.000 | 3.266 | -2.266 | 1.721 | 1.317 | 4.000 | 0.459692 |
| ADAL | No | 0.253394 | 1.000 | 3.751 | -2.751 | 2.063 | 1.333 | 4.000 | 0.459692 |
| AICDA | No | 0.310537 | 1.000 | 2.151 | -1.151 | 0.9918 | 1.160 | 4.000 | 0.459692 |
| DPP4 | No | 0.189521 | 1.000 | 1.375 | -0.3751 | 0.2376 | 1.579 | 4.000 | 0.438131 |
| FOLR4 | No | 0.274663 | 1.000 | 2.239 | -1.239 | 0.9797 | 1.265 | 4.000 | 0.459692 |
| ICAM1 | No | 0.404114 | 1.000 | 2.404 | -1.404 | 1.506 | 0.9320 | 4.000 | 0.459692 |
| IL19 | No | 0.287643 | 1.000 | 2.799 | -1.799 | 1.468 | 1.225 | 4.000 | 0.459692 |
| IL25 | No | 0.357345 | 1.000 | 2.257 | -1.257 | 1.210 | 1.039 | 4.000 | 0.459692 |
| IL33* | No | 0.012223 | 1.000 | 2.068 | -1.068 | 0.2459 | 4.343 | 4.000 | 0.129937 |
| IL4 | No | 0.401930 | 1.000 | 2.333 | -1.333 | 1.423 | 0.9367 | 4.000 | 0.459692 |
| ITGA2B | No | 0.299923 | 1.000 | 3.294 | -2.294 | 1.928 | 1.190 | 4.000 | 0.459692 |
| ITGA6 | No | 0.015643 | 1.000 | 2.073 | -1.073 | 0.2657 | 4.037 | 4.000 | 0.147938 |
| LTBR** | No | 0.006280 | 1.000 | 2.344 | -1.344 | 0.2559 | 5.254 | 4.000 | 0.129937 |
| Pdgfb | No | 0.075183 | 1.000 | 2.073 | -1.073 | 0.4488 | 2.390 | 4.000 | 0.307221 |
| ZBTB7B | No | 0.036957 | 1.000 | 2.417 | -1.417 | 0.4603 | 3.079 | 4.000 | 0.217268 |
